# Deep-sea fish reveal alternative pathway for vertebrate visual development

**DOI:** 10.1101/2024.10.10.617579

**Authors:** Lily G. Fogg, Stamatina Isari, Jonathan E. Barnes, Jagdish Suresh Patel, N. Justin Marshall, Walter Salzburger, Fabio Cortesi, Fanny de Busserolles

## Abstract

Vertebrate vision is accomplished by two phenotypically distinct types of photoreceptors in the retina: the saturation-resistant cones for the detection of bright light and the highly sensitive rods for dim light conditions [1]. The current dogma is that, during development, all vertebrates initially feature a cone-dominated retina, and rods are added later [2, 3]. By studying the ontogeny of vision in three species of deep-sea fishes, we show that their larvae express cone-specific genes in photoreceptors with rod-like morphologies. Through development, these fishes either retain this rod-like cone retina (*Maurolicus mucronatus*) or switch to a retina with true rod photoreceptors with expression of rod-specific genes and transcription factors (*Vinciguerria mabahiss* and *Benthosema pterotum*). In contrast to the larvae of most marine fishes, which inhabit the bright upper layer of the open ocean, the larvae of deep-sea fishes occur deeper, exposing them to a dimmer light environment [4–7]. Spectral maxima predictions from molecular dynamics simulations and environmental light estimations suggest that using transmuted photoreceptors that combine the characteristics of both cones and rods maximises visual performance in these dimmer light conditions. Our findings provide molecular, morphological, and functional evidence for the evolution of an alternative developmental pathway for vertebrate vision.

## Introduction

Vertebrate vision has evolved to function in diverse photic environments, from bright and colourful ecosystems, such as coral reefs and rainforests, to the near darkness found in caves and the deep sea. In the vast majority of vertebrates, vision is accomplished by the interplay of two types of retinal photoreceptors: rods and cones [8]. Rods are characterized by a specialized morphology tailored towards photon capture with densely packed visual pigments (opsins) and a highly sensitive phototransduction pathway, and function in dim-light (scotopic) conditions (Table 1). In contrast, cones are morphologically and molecularly primed for bright-light (photopic) conditions [1]. In intermediate (mesopic) light conditions, rods and cones usually work together [9]. In a few species, however, transmuted (“hybrid”) photoreceptors have been documented, which combine features of both cell types. These taxa typically inhabit mesopic light environments (*e.g.,* pearlside fishes [10], lampreys [10], and skates [11]) or have experienced a switch in diel activity period (*e.g.,* snakes and geckos [11]).

**Table 1.** Summary of the characteristics of rod-like cones of different species compared to true rods and true cones. Adapted from de Busserolles et al., 2017 [10]. *, this study. Salamander is the tiger salamander, *Ambystoma tigrinum*. Pearlside is *Maurolicus muelleri* (adults) or *M. mucronatus* (larvae). Snake data are for the nocturnal genus *Hypsiglena*. The gecko is the nocturnal Tokay gecko, *Gekko gekko*. Lightfish is *Vinciguerria mabahiss.* Lanternfish is the Skinnycheek lanternfish, *Benthosema pterotum*. R, rod-like; C, cone-like; n.a., not available; poly, polysynaptic.

|  |  |  | Rod-like cone |  |  |  |  |  |  |
| --- | --- | --- | --- | --- | --- | --- | --- | --- | --- |
| Photoreceptor Characteristics | True rod [1, 73] | True cone [1, 73] | Gecko [53, 54, 56] | Snake [51, 74] | Salamander [49, 50] | Pearlside [10] | Pearlside larvae (*) | Lightfish larvae (*) | Lanternfish larvae (*) |
| Outer segment shape | Long, rod-shaped (cylindrical) | Short, cone-shaped (distally tapering) | R | R | R | R | R | R | R |
| Outer segment discs | Individual sealed disc, separated from plasma membrane | Disks continuous with plasma membrane (open) | R C | n.a. | n.a. | R | R | R | R |
| Incisure | Present | Absent | R | n.a. | R | R | C | C | C |
| Paraboloid | Absent | Present | C | R | R | R | R | R | R |
| Oil droplet | Absent | Sometimes | R | R | R | R | R | R | R |
| Synaptic ending | Small, spherical, oligosynaptic | Large, conical, flat-end base, polysynaptic | C | n.a. | R C<br>Small poly | R C<br>Small poly | n.a. | n.a. | n.a. |
| Opsin | *rh1* | *sws1*, *sws2*, *lws*,<br>*rh2* | C<br>*rh2* | C<br>*sws1*<br>*lws* | C<br>*sws2* | C<br>*rh2-1*<br>*rh2-2* | C<br>*rh2-1*<br>*rh2-2* | C<br>*rh2* | C<br>*rh2* |
| Spectral sensitivity (nm) | 480-510 nm | *rh2*, 450–530<br>*sws1*, 360–440<br>*sws2*, 400–450<br>*m/lws*, 510–560 | C<br>521 | C<br>358,<br>536 | C<br>432 | C<br>441<br>(both; [10]);<br>474,<br>480 (*) | C<br>474,<br>480 | C<br>474 | C<br>470 |
| Phototransduction cascade | Rod-like<br>(*e.g., gnat1, saga, sagb*) | Cone-like<br>(*e.g., gnat2, arr3a, arr3b*) | C(R) | n.a. | R | C | C | C | C |
| Cell physiology | Rod properties<br>(high sensitivity) | Cone properties<br>(fast, never saturate) | R | n.a. | n.a. | n.a. | n.a. | n.a. | n.a. |

The balance between rods and cones in the retina is finely tuned to ecological demands [12–15]. For example, the spectral sensitivity of each photoreceptor type is typically tuned to the wavelength of the prevailing light via their opsin pigments. The rod opsin (RH1) is sensitive to a narrow range of blue-green light and multiple cone opsins (SWS1, SWS2, RH2 and LWS) cover a broad sensitivity range from ultraviolet to red [16]. Furthermore, while diurnal vertebrates typically have higher cone densities, their nocturnal counterparts tend to have rod-dominated retinas [17–19]. This is pushed to the extreme in species with pure rod retinas, such as some nocturnal reptiles [20] and many deep-sea fishes [21]. Nevertheless, the developmental trajectories that give rise to this retinal diversity are remarkably conserved: vertebrates start their lives with cone-dominated retinas, with rods emerging later in ontogeny [2, 3]. This “cones first – rods later” pathway of retinal development suits the ecology of terrestrial vertebrates and most marine fishes, which initially inhabit the bright upper pelagic ocean. However, it is seemingly mismatched with the lifestyle of deep-sea fishes that spend their entire lives in deeper and dimmer waters [4–7], and are characterized by morphologically rod-dominated retinas from early developmental stages onwards [2, 22–24].

Like many other facets of their biology, our understanding of the development of the visual system of deep-sea fishes is limited and, in part, contradictory. We thus set out to examine in detail the retinal development of three deep-sea fish species: the lightfish *Vinciguerria mabahiss* (Stomiiformes: Phosichthyidae), the hatchetfish *Maurolicus mucronatus* (Stomiiformes: Sternoptychidae), and the lanternfish *Benthosema pterotum* (Myctophiformes: Myctophidae). These three species reside in different photic niches, offering a unique opportunity to study photoreceptor development. Between larval and adult stages, *V. mabahiss* switches from mesopic to scotopic conditions, *B. pterotum* switches from photopic-mesopic to scotopic conditions, and *M. mucronatus* remains in mesopic conditions throughout life [4, 6, 25–29]. Using light and electron microscopy, bulk retinal transcriptome sequencing, amino acid sequence analysis, and spectral sensitivity predictions, we bridge the gap between gene expression and morphology in deep-sea fishes to reveal how vision develops in one of the dimmest and largest habitats on Earth.

## Results and Discussion

### Visual gene expression

The analysis of bulk retinal transcriptomes revealed that early larval stages of all three species expressed exclusively or predominantly the green-sensitive cone opsin, *rh2,* and the cone-specific phototransduction genes*, gnat2*, *arr3a* and *arr3b* (Fig. 1; Table S2). This is congruent with findings in other deep-sea fish larvae [30]. Conversely, no expression of the rod opsin (*rh1*) or rod-specific phototransduction genes (*gnat1*, *saga* and *sagb*) was observed in the early stages of *V. mabahiss*. Low levels of *rh1* but no other rod-related phototransduction genes were detected in larval *B. pterotum* (0.2% of total opsin expression at pre-flexion, 3% at post-flexion), and low expression levels of all rod-specific genes were found in larvae of *M. mucronatus* [*rh1* (0.2%), *gnat1* (0.1%), *saga* and *sagb* (0.3%) at post-flexion]. Like in other deep-sea fishes [30], an ontogenetic switch from predominantly cone-specific to rod-specific visual gene expression was observed in *V. mabahiss* and *B. pterotum* (Fig. 1; Fig. S1-4; Table S2). For the species sampled at higher temporal density during development (*V. mabahiss*), we could narrow down the timing of this switch to metamorphosis (*i.e.,* between mid and late post-flexion), coinciding with the onset of this species’ migration to deeper and dimmer waters [4]. In contrast, *M. mucronatus* continued to predominantly express cone opsin genes into adulthood [*rh2* (99.6%), cone arrestins (98.2%), and cone transducins (99.7%)], similar to previous findings for *M. muelleri* [10]. Hence, molecularly, the vertebrate cone-to-rod pathway is conserved in *V. mabahiss* and *B. pterotum,* but not in *M. mucronatus*.

**Fig. 1.**
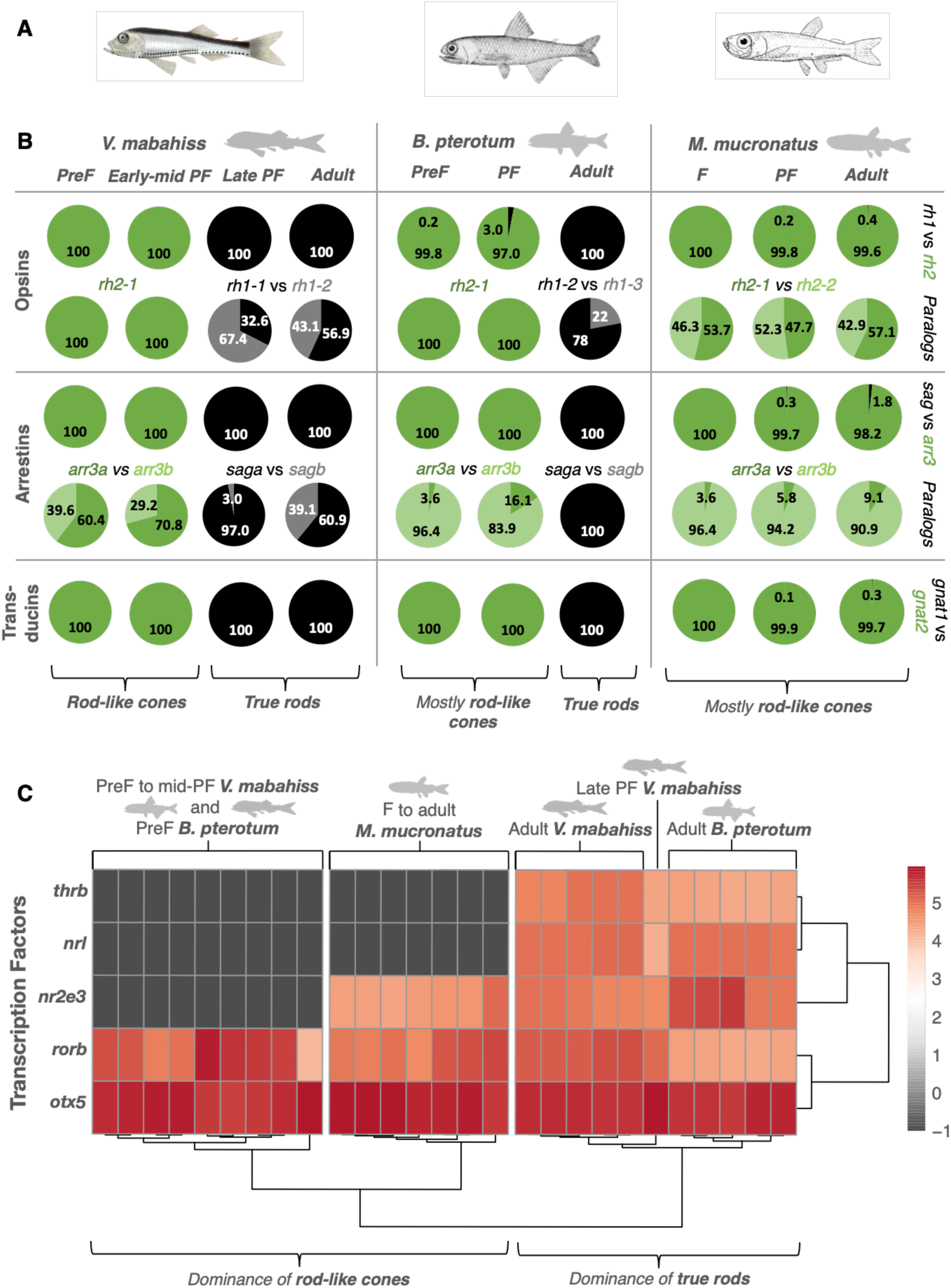
Molecular basis for visual development in deep-sea fishes. **A.** Illustrations of the species used in this study (left to right): *Vinciguerria mabahiss* [75], *Benthosema pterotum* [76] and *Maurolicus mucronatus* [77]. **B.** Mean proportional expression of opsin, arrestin and transducin genes in the retina over ontogeny (given as a % of total gene family expression for subclasses or as a % of total gene subclass expression for paralogs). Cone-specific genes are coloured green while rod-specific genes are coloured grey/black. **C.** Heatmap showing per-specimen retinal expression (in log_10_TPM) of transcription factors involved in photoreceptor development. PreF, pre-flexion; F, flexion; PF, post-flexion. *arr3*, arrestin 3; *sag*, S-antigen arrestin; *rh1*, rhodopsin 1 (rod opsin); *rh2*, rhodopsin 2; *otx5,* orthodenticle homolog 5*; rorb,* RAR related orphan receptor B; *nr2e3,* nuclear receptor subfamily 2 group E member 3*; thrb,* thyroid hormone receptor β*; nrl,* neural retina leucine zipper.

Detailed transcriptome mining and phylogenetic reconstructions (Fig. S1-2) revealed that two *rh2* paralogs were expressed in *M. mucronatus* throughout life. In contrast, a single *rh2* was expressed in larval *V. mabahiss* and *B. pterotum* (Table S2). The inverse was true of *rh1*, with two and three different paralogs expressed in adult *V. mabahiss* and *B. pterotum,* respectively, but only one in *M. mucronatus* throughout life. Several *rh2* paralogs are common in many teleost species, especially in deep-sea fishes [31]. In contrast, most vertebrates have only one *rh1* [32], and only a handful of teleosts (mainly deep-sea species) have been shown to express multiple paralogs, including another species in the genus *Benthosema* [33]. The expression of several opsin genes suggests the presence of photoreceptors with distinct spectral sensitivities in all species.

### Spectral sensitivities

Using spectral maxima predictions based on an experimentally validated molecular dynamics simulations-based approach [33, 34], we found that all three species examined had at least two spectrally distinct visual opsins over ontogeny (Fig. 1B). Specifically, *V. mabahiss* and *B. pterotum* each had one RH2 (sensitive to 474 nm and 470 nm, respectively) as larvae, but switched to two RH1 pigments (sensitive to wavelengths between 496-499 nm in *V. mabahiss* and 498-504 nm in *B. pterotum*) as adults. Conversely, *M. mucronatus* had the same dominant visual opsins throughout life (two RH2s covering a 474-481 nm range). These spectral sensitivities correlate with the light environment in which the different developmental stages occur [6, 10, 25, 28]. We thus demonstrate that, just like in shallow-water fishes [13, 16, 35], spectral tuning during ontogeny matches the prevailing light environment in deep-sea fish species. The expression of multiple visual opsin genes with distinct spectral sensitivities in all three species under investigation further suggests the presence of several morphologically distinct photoreceptor types in their retinas.

### Photoreceptor Morphology

Most vertebrates have a duplex retina containing both rods and cones, while many deep-sea fishes feature pure rod retinas, at least as adults [21]. Based on light and electron microscopy, we found a dominance of morphologically rod-like photoreceptors in the early developmental stages of all three study species (Fig. 2; Fig. S5). All photoreceptors examined in the larvae of *V. mabahiss* and *M. mucronatus* had long and cylindrical outer segments (OS), closed OS disc membranes, and lacked a paraboloid or oil droplet, all of which are anatomical features typical of rods. Similarly, most photoreceptors in larval *B. pterotum* were rod-like (Fig. S5); however, a few cone-shaped cells with short, tapered OS and open, continuous OS disc membranes were also observed (<10% of photoreceptors examined) (Fig. S6). This is in line with previous work documenting morphologically rod-like cells in the larvae of other deep-sea fishes, including lanternfishes [23, 26], a Macrouridae species [2], *Evermannella sp., Paraliparis sp.* and *Idiacanthus fasciola* [22]. However, the incidence of exclusively (in *V. mabahiss* and *M. mucronatus*) or predominantly (in *B. pterotum*) morphologically rod-like photoreceptors coincided with predominantly cone-specific retinal gene expression patterns (≥97%). This indicates that the retina is dominated by transmuted photoreceptors and strongly suggests that the three deep-sea fish species studied here diverge from the conserved cone-to-rod pathway of other vertebrates.

**Fig. 2.**
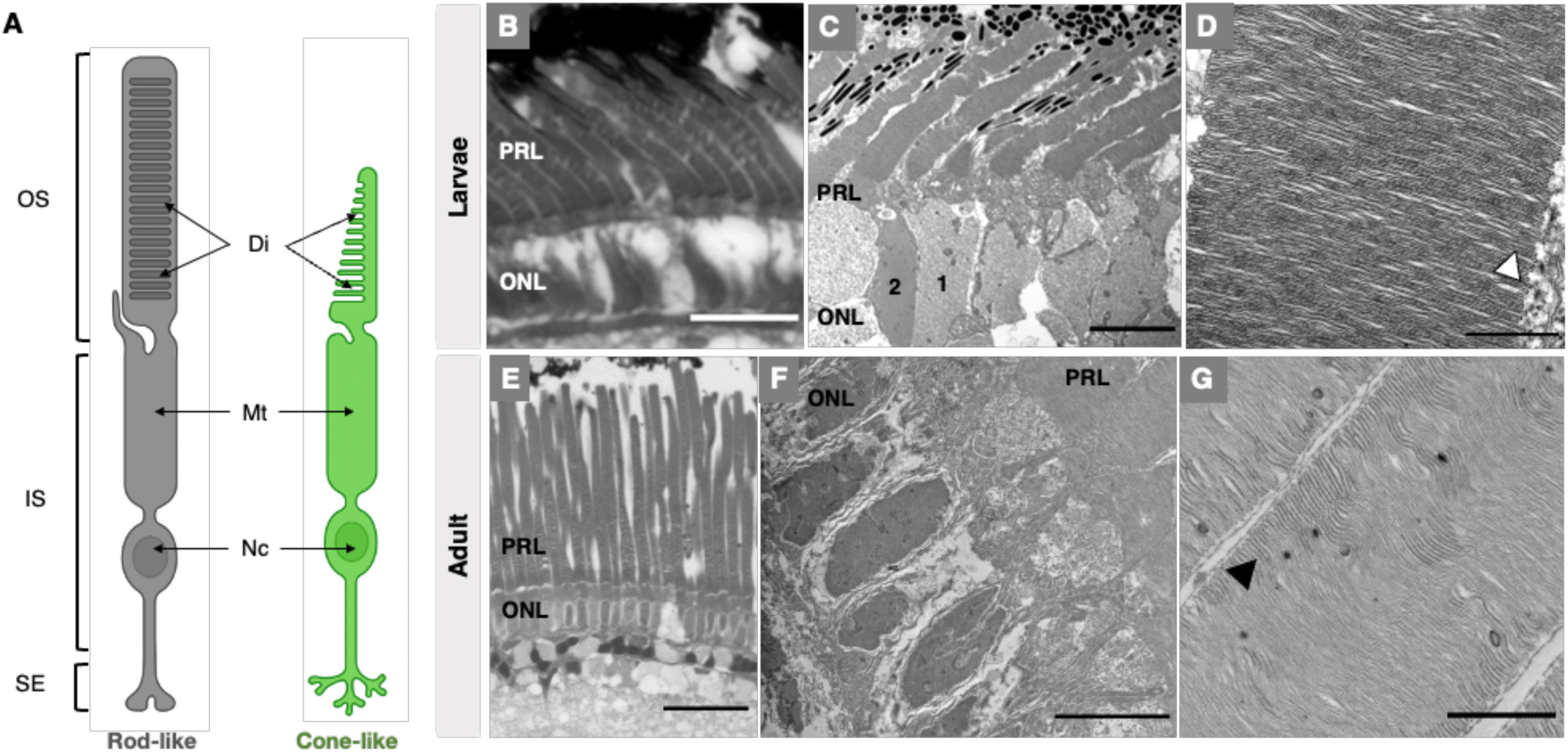
Morphological basis for visual development in deep-sea fishes. **A.** Schematic of photoreceptors with rod-like or cone-like morphology**. B-G.** Representative light (**B, E**) and transmission electron (**C-D, F-G**) micrographs of the photoreceptor layers in early larval and adult *V. mabahiss*. In both larvae and adults, the retina was dominated by morphologically rod-like photoreceptors, with long, cylindrical outer segments (**B-C, E-F**) and closed outer segment discs (**D, G**; arrowhead). Notably, larvae had two morphological types of nuclei in the outer nuclear layer (ONL) characterised by lighter (type 1) or darker (type 2) chromatin staining (**C)**, while adults had only one type with darker chromatin staining (**F**). OS, outer segment; IS, inner segment; SE, synaptic ending; Di, discs; Mt, mitochondria; Nc, nucleus; PRL, photoreceptor layer; ONL, outer nuclear layer. Scale bars: B, 10 µm; C, F, 5 µm; D, 500 nm; E, 25 µm; G, 1 µm.

The adults of all three species had purely morphologically rod-like photoreceptors, which coincided with rod-specific gene expression in *V. mabahiss* and *B. pterotum* and cone-specific gene expression in *M. mucronatus* (Fig. 1-2; Fig. S5). This indicates an adult retina composed purely of true rods in *V. mabahiss* and *B. pterotum*, similar to many other deep-sea fishes [21]. However, as shown before [10], *Maurolicus* spp. have adult retinas dominated by rod-like cones (99-99.6%), with a small population of true rods (0.4-1%) (Fig. S5). While the morphology of larval and adult photoreceptors was quite similar, the photoreceptors of adults of all species had substantially longer OS as well as incisures (Fig. S7). Furthermore, larvae of all species had two morphologically distinct types of photoreceptor nuclei, while the adults of *V. mabahiss* and *B. pterotum* had only one type. In larvae, lighter-staining nuclei dominated, congruent with the cone expression data, while the inverse was true of adults in which rod gene expression dominated.

### Developmental transcription factor expression

Previous work has shown that transcription factors (TFs), such as OTX5 (known as CRX in mammals), RORβ, NR2e3, NRL, and THRβ play a coordinated role in directing retinal progenitor cells towards either a true cone or true rod cell fate. However, the regulation of transmuted photoreceptor development is unknown [36]. We thus examined TF expression in the three deep-sea species to understand the regulatory factors governing the development of transmuted photoreceptors. We uncovered that *otx5* and *rorβ* were consistently expressed in all species at stages with rod-like cones, including earlier stages of *V. mabahiss* and *B. pterotum* and all stages of *M. mucronatus* (Fig. 1; Table S3). Notably, these TFs continued to be expressed in adults with only true rods (*V. mabahiss* and *B. pterotum*). This suggests that OTX5, a TF associated with both rod and cone development in zebrafish [37, 38], and RORβ, a rod-associated TF in mice [39], may direct rod-like cone development in larvae and true-rod development later in ontogeny.

In mammals, the synergistic action of OTX5 and RORβ activates the expression of short-wavelength opsin genes [40]. Interestingly, mammals recruit rods from a short-wavelength-sensitive cone lineage during ontogeny, a likely remnant of the selective pressures experienced when switching to nocturnality during the Mesozoic [41, 42]. Deep-sea fishes may experience comparable selective pressures due to migration from bright shallow waters to the extremely dim deep sea. Thus, they may have convergently evolved or retained an ancestral pathway that redirects a cone fate to rod-like cones in larvae before producing true rods in adults. Finally, since the use of rod-like cones may facilitate the transition to a pure rod retina both ontogenetically and evolutionarily, this adaptation may even have been important during the colonisation of the deep sea.

We also found that the expression of *nr2e3,* a rod-specific TF in vertebrates [43–45], correlates with the presence of true rods in deep-sea fishes. Furthermore, *B. pterotum* did not express *nr2e3* at earlier stages when its true rods were not likely to be functional yet (*i.e.,* rod opsin and rod-like morphology were present, but rod phototransduction gene expression was absent; Fig. 1). Hence, we propose a temporal association between *nr2e3* expression and the functional maturation of true rods in these species.

*Nrl* and *thrβ* were associated with a dominance of true rods in adults of *B. pterotum* and *V. mabahiss*. Co-expression of these TFs can produce rods in mice [46]. Notably, *nrl* was absent in late post-flexion *V. mabahiss*, which likely has an immature true rod retina (Fig. 1). Therefore, NRL may be restricted to rod specification or maintenance in adult deep-sea fishes and dispensable earlier in ontogeny, similar to adult mammals [47]. This further supports that deep-sea fishes utilise an alternative, NRL-independent pathway to specify rod fate earlier in ontogeny, similar to Atlantic cod [48].

### Photoreceptor transmutation in larval deep-sea fishes

To the best of our knowledge, our study is the first to report the discovery of larval retinas dominated by rod-like cones in vertebrates. The only other vertebrate known to have transmuted photoreceptors as larvae is the tiger salamander (*Ambystoma tigrinum*), in which rod-like cones represent ≤ 1% of photoreceptors [49, 50]. In contrast, rod-like cones were already dominant in the youngest specimens in our study, which were sampled shortly after hatching as pre-flexion larvae (Fig. 1-2). Possessing rod-like cones as larvae may make the transition to the pure rod retina of adults more rapid and metabolically efficient. Furthermore, since vision is primarily used after hatching, it is very likely that rod-like cones are the first functionally relevant photoreceptors in these fishes. This is well aligned with ecology of these species, as combining photoreceptor characteristics into a single rod-like cone is likely the most efficient way to optimise vision in the mesopic conditions which these fish experience (Fig. 3) [10]. Notably, transmuted photoreceptors were originally proposed to be an evolutionary intermediate between the two canonical photoreceptor types, rods and cones, that arose after an ecological shift [51]. However, since transmuted photoreceptors are well-suited to the ecology of larval deep-sea fishes, it is more likely in this case that they represent a newly evolved photoreceptor type adapted for a mesopic photic environment. Further work is required to determine whether transmuted photoreceptors should be re-classified as a novel photoreceptor type, rather than a “hybrid” intermediate between rods and cones.

**Fig. 3.**
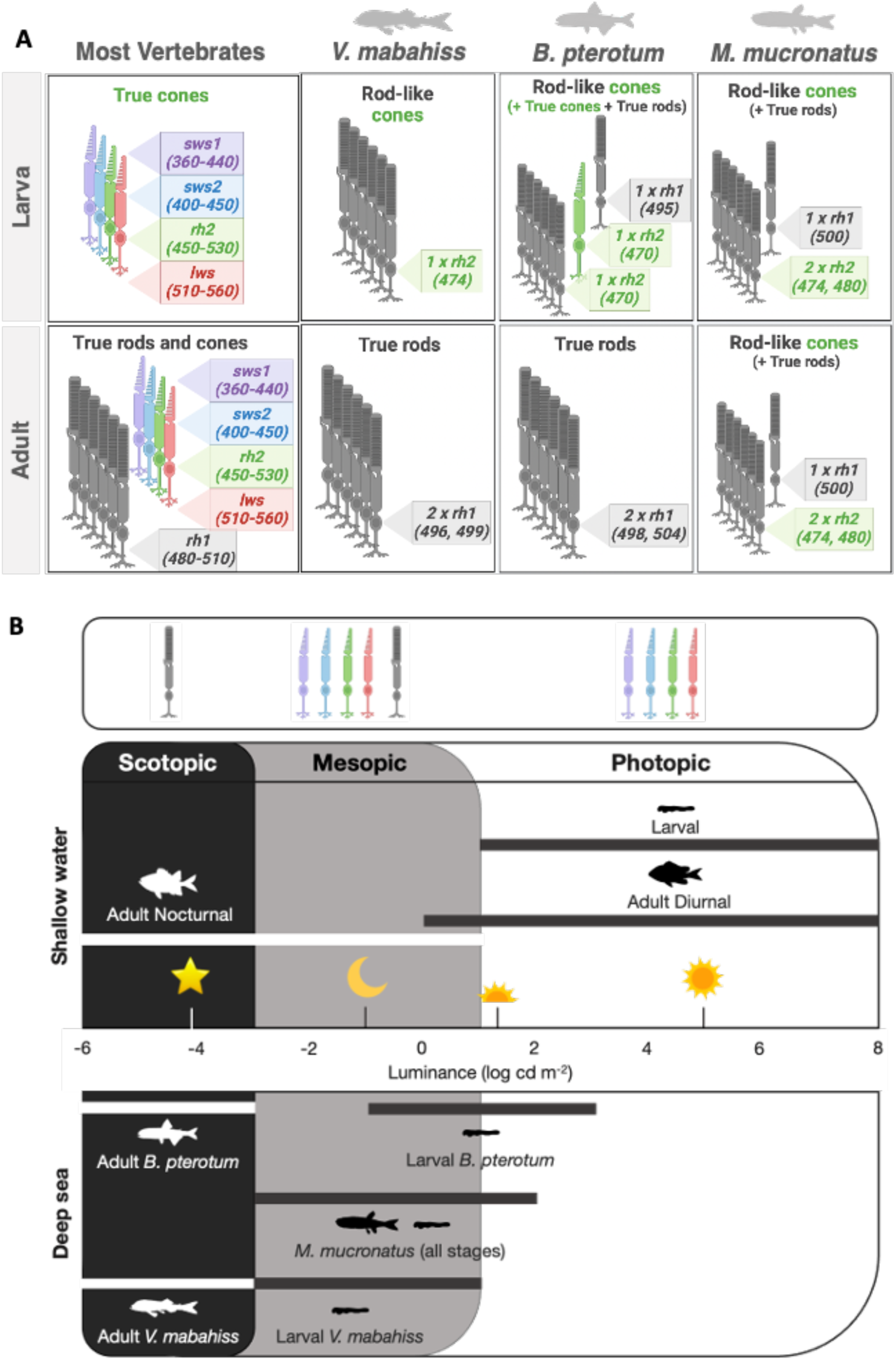
Functional relevance of novel developmental pathway in deep-sea fishes. **A.** Schematic of photoreceptor development showing the conserved cone-to-rod pathway of most vertebrates and the developmental models for the deep-sea fish species from this study. Photoreceptor cartoon denotes morphology (rods coloured grey, cones coloured green), while opsin gene expression (subclass, copy number and predicted spectral maxima) is detailed in inset boxes. Note that larval deep-sea fish possess mostly rod-like cones, while most vertebrates have true cones as larvae. **B.** Schematic of photic environments of shallow-water and deep-sea fishes over ontogeny (coloured boxes), showing the functional ranges of different photoreceptor types (arrows) and light environments experienced by different species and life stages (inset bars) [6, 10, 25, 28]. In shallow-water fish, the larvae inhabit photopic conditions and the retina has only true cones. Conversely, adults inhabit dimmer conditions, and the retina has both true rods and true cones. In deep-sea species, larvae inhabit predominantly mesopic conditions and have mostly rod-like cones. In adults, *M. mucronatus* remains in mesopic conditions and retains rod-like cones, while the other two species migrate to scotopic conditions and adopt pure rod retinas. Environmental light sources (from left to right) are as follows: starlight, full moon, civil twilight, sunset/sunrise, and sunlight [78]. Figure partially redrawn from de Busserolles et al., 2017 [10].

Our data also suggest that photoreceptor transmutation in vertebrates may be more widespread than previously thought. The fact that the species in this study fall into two phylogenetically distant clades [Stomiati (*V. mabahiss* and *M. mucronatus*) and Neoteleostei (*B. pterotum*)] that diverged nearly 200 Mya [52] combined with the shared mesopic conditions of many deep-sea larvae, suggests that transmutation could be much more common across the deep-sea teleost phylogeny. This is further supported by at least another seven deep-sea species that are known to possess morphologically rod-like photoreceptors as larvae [2, 22, 23, 26], and at least eleven other species predominantly expressing cone opsin genes early in development [30]. Moreover, although the current study is the first to examine morphology and gene expression together, two other studies independently showed rod-like photoreceptors [22] and cone opsin gene expression [30] in larval stages of another deep-sea fish: *Idiacanthus fasciola*.

Finally, photoreceptor transmutation has also been reported in taxa beyond the ray-finned fishes, including reptiles (geckos [53–56] and snakes [57]), amphibians (salamanders [49, 50]), cartilaginous fish (skate [58–60]), and jawless fish (lampreys [61, 62]). The distribution of these photoreceptors across most vertebrate classes, as well as their presence in an early diverging lineage (Agnatha), suggests that they could have evolved early on during the diversification of vertebrates. Further work on species which experience mesopic conditions, such as crepuscular species, will be required to determine the prevalence of transmuted photoreceptors across the vertebrate phylogeny and to explore the evolutionary history of this potentially novel photoreceptor type.

### Conclusion

Using light and electron microscopy, bulk transcriptome sequencing, amino acid sequence analysis, and spectral sensitivity predictions, we reveal a novel pathway for vertebrate visual development in deep-sea fishes. We found that several phylogenetically distant deep-sea fishes have retinas dominated by transmuted photoreceptors at early life stages, combining the molecular machinery of cones with the morphology of rods to generate rod-like cones. These transmuted photoreceptors are retained through to adulthood in *M. mucronatus*, while *B. pterotum* and *V. mabahiss* later adopted retinas dominated by true rods. These photoreceptor types are well suited to the photic conditions at each life stage. Furthermore, we identified candidate TFs involved in the development of transmuted and true photoreceptors in deep-sea fishes. Our findings advance our understanding of the evolutionary dynamics of vision in unconventional and extreme environments and challenge the existing paradigms for the classification of photoreceptors and the development of vision in vertebrates.

## Acknowledgements

We would like to thank the staff of the research vessels, the Thuwal and the R/V Al Azizi, for support during field work. We thank Robert Sullivan from the Queensland Brain Institute (QBI) Histology Facility, Richard Webb and Robyn Chapman Webb from the Centre for Microscopy and Microanalysis (CMM) and Rumelo Amor from the QBI Advanced Microscopy facility for technical support and advice. We acknowledge the staff at Novogene Co., Ltd for library preparation and transcriptome sequencing. We are grateful to Alexander Davis from Duke University for the generous contribution of samples. We acknowledge Research Computing and Data Services in the Institute for Interdisciplinary Data Science at the University of Idaho for computational resources for molecular dynamics simulations. Finally, we acknowledge that panels in some figures were created using BioRender.

## Data Availability

Newly identified sequences and sequenced transcriptomes will be available through GenBank and the SRA archive upon publication. All other data will be available via Dryad (DOI) upon publication or are provided in the main manuscript or Supplemental Information.

## Methods

### Animal collection and tissue preservation

Three species of deep-sea fishes were collected for this study: *V. mabahiss, B. pterotum* and *M. mucronatus* (Table S1). All specimens were collected from the Saudi Arabian Red Sea. Larvae were collected aboard the research vessel R/V Al Azizi in March 2018. Larvae were sampled at 0-200 metres depth during the day and night using oblique bongo net tows. Adults were collected aboard the research vessel Thuwal in July 2014 using a Tucker trawl.

Following collection, fishes were sorted and identified on board to the lowest taxonomic level possible. For each species, individuals were pooled by size and/or developmental stage and fixed whole in either 4% paraformaldehyde [PFA; 4% (w/v) PFA in 0.01M phosphate-buffered saline] or RNAlater. Following fixation, fish were imaged alongside a scale reference under a dissection microscope and eyes were enucleated for processing. The standard length was subsequently measured from images using Fiji v1.53c [63]. All procedures were approved by the University of Queensland Animal Ethics Committee (ANRFA/014/18). The animal collection was in accordance with the regulations of the King Abdullah University of Science and Technology, Saudi Arabia.

### Transcriptome sequencing, quality control and de novo assembly

Retinal transcriptomes were sequenced for a total of 21 individuals: *V. mabahiss* [pre-flexion larvae (*n*=3), early-mid post-flexion larvae (*n*=4), late post-flexion larvae (*n*=1), adults (*n*=5)], *B. pterotum* [pre-flexion larvae (*n*=1), post-flexion larvae (*n*=1), adults (*n*=4)] and *M. mucronatus* [flexion larvae (*n*=1), post-flexion larvae (*n*=1)]. The adult dataset was completed with previously published transcriptomes [*B. pterotum* (*n*=1), *M. mucronatus* (*n*=5)] [10, 33], resulting in a total dataset of 27 retinal transcriptomes spanning several life stages in each of the species.

For all samples, retinal tissue was digested with Proteinase K (New England Biolabs) for 15-30 min at 55°C. Total RNA was extracted and isolated using the Arcturus PicoPure RNA Isolation Kit (Applied Biosystems) and the Monarch Total RNA Miniprep Kit for larvae and adults, respectively. Genomic DNA was removed from all samples with RNase-free DNase, and the quality and yield of isolated RNA were assessed using Eukaryotic Total RNA 6000 kits (Pico kit for larvae and Nano kit for adults; Agilent Technologies) on the Queensland Brain Institute’s Bioanalyser 2100.

RNA extractions were shipped on dry ice and whole-retina transcriptome libraries were prepared from total RNA at Novogene’s sequencing facilities in Hong Kong and Singapore. The Clontech SMART-Seq v4 Ultra Low Input RNA Kit and the NEBNext Ultra RNA Library Prep Kit for Illumina were used for larval samples. The NEBNext Ultra RNA library preparation kit for Illumina was used for adult samples. The concentration of each library was checked using a Qubit dsDNA BR Assay kit prior to barcoding and pooling at equimolar ratios. Libraries were sequenced as 150 bp paired-end reads on an Illumina NovaSeq 6000 S4 flow cell. Libraries were trimmed and *de novo* assembled as per the methods described in de Busserolles et al., 2017 [10]. Briefly, read quality was assessed using FastQC (v0.72), raw reads were trimmed and filtered using Trimmomatic (v0.36.6) and transcriptomes were *de novo* assembled with Trinity (v2.8.4) using the genomics virtual laboratory on the Galaxy platform at usegalaxy.org [64].

### Visual gene mining and differential expression analyses

Published cytochrome C oxidase subunit I (*COI*), opsin, transducin and arrestin gene sequences for *M. mucronatus* were obtained from GenBank. The remaining COI, opsin, transducin, arrestin and transcription factor (TF) genes were mined from the transcriptome in Geneious Prime v2021.1.1 (Biomatters Ltd, version 2019.0.4). All expression analyses were also conducted in Geneious Prime. Initially, *COI* genes were extracted from *de novo* assembled transcriptomes for species identification by mapping to species-specific references from Genbank (https://www.ncbi.nlm.nih.gov/genbank/) with medium sensitivity settings. Opsin, transducin, arrestin and TF gene extractions were performed by mapping assembled transcriptomes to published coding sequences (CDS) for the most phylogenetically similar species available on Genbank using customised sensitivity settings (fine-tuning, none; maximum gap per read, 15%; word length, 14; maximum mismatches per read, 40%; maximum gap size, 50 bp; index word length, 12; paired reads must both map). Contigs mapped to references were scored for similarity against publicly available sequences using BLASTn (NCBI, Bethesda, MD, https://blast.ncbi.nlm.nih.gov/Blast.cgi). One of the limitations of *de novo* assembly of highly similar genes (such as opsin gene paralogs) using short-read transcripts is that it can produce erroneous (chimeric) sequences or fail to reconstruct lowly expressed transcripts. Thus, for the opsin genes, a second approach was also employed using a manual extraction method from back-mapped reads to verify the initially extracted opsin genes, as per de Busserolles et al., 2017 [10].

During manual gene extraction, filtered paired reads were mapped against the closest reference CDS (with previously stated customised sensitivity settings). Matching reads were connected by following single nucleotide polymorphisms (SNPs) across genes with continual visual inspection for ambiguity and were extracted as paired mates to mitigate sequence gaps. The consensus of an assembly of these extracted reads was used as the reference for low sensitivity (high accuracy, 100% identity threshold) mapping. Partial CDS extractions were cyclically mapped using the low-sensitivity approach to prolong and subsequently remap reads until a complete CDS was obtained.

To confirm the identity of all genes mined from the transcriptome, full coding sequences were checked on BLASTn. Subsequently, opsin, transducin and arrestin genes were further characterised using gene phylogenies. Briefly, the extracted CDS were used in conjunction with reference datasets obtained from Genbank (www.ncbi.nlm.nih.gov/genbank/) and Ensembl (www.ensembl.org/) to phylogenetically reconstruct separate gene phylogenies [10]. All gene sequences were aligned using the MUSCLE plugin v3.8.425 [65] in Geneious Prime. MrBayes v3.2.6 [66] on CIPRES [67] was used to reconstruct phylogenetic trees from the aligned sequences using the following parameters for each reconstruction: GTR+I+G model, two independent MCMC searches with four chains each, 10 million generations per run, 1000 generations sample frequency, and 25% burn-in. The generated trees were manually edited in Figtree v1.4.4 [68].

For differential expression analyses, gene paralogs were first scored on similarity using pairwise/multiple alignments. The similarity score minus one was used as the gene-specific maximum % mismatch threshold for mapping (paired) transcripts back onto complete extracted opsin CDS to ensure that reads did not map to multiple paralogs. Absolute gene expression in log_10_TPM (Transcripts Per Kilobase Million) was then calculated as follows, where *T_gene_* represents the number of transcripts mapped to each gene and *T_transcriptome_*, is the number of transcripts per transcriptome:

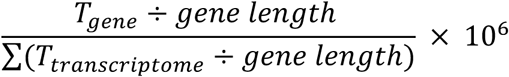

Finally, proportional gene expression was also determined for the opsin, transducin and arrestin genes. Proportional expression was calculated as the number of reads mapped to a particular gene (*e.g., rh1*) divided by the number of reads mapped to all genes in that family (*e.g.,* all opsins), adjusted to account for differing gene lengths. Further proportional comparisons were made within subfamilies using the same method, for example, to find the proportional expression of a particular paralog (*e.g., rh1-1*) within a subfamily (*e.g., rh1*).

### Spectral maxima predictions based on atomistic molecular dynamics simulations

Opsin gene sequences mined from the transcriptomes were translated and used to determine the peak spectral sensitivities (λ_max_) of each of the 10 deep-sea visual photopigments using experimentally validated statistical models based on dynamical features derived from the molecular dynamics simulations [33, 34, 69]. The statistical model used for RH1 opsins was:

λ_max_ = 831.762 + 5.851 × Angle 3 - 2.997 × Torsion 15 + 45.585 × DisulfideBridge While for RH2 opsins the model was:

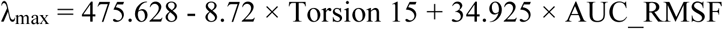

Here, Angle 3 (C3–C7–C8) and Torsion 15 (C7–C6–C5–C18) are the median values obtained from the molecular dynamics simulations, AUC_RMSF is the area under the curve (AUC) of the root mean square fluctuations (RMSF) of the chromophore 11-*cis* retinal bound to known lysine residue, and DisulfideBridge indicates the presence or absence of a cystein-cystein bridge between aligned positions 111 and 188 in the opsin protein.

The opsin amino acid sequences from 10 deep-sea visual photopigments were used as input to calculate the parameters needed for these statistical models. Structures were prepared for all opsin sequences using Alphafold v2.3.1 [70] with GPU relaxation. The top minimized 3-dimensional structure (measured by the predicted local distance difference test - pLDDT) for each deep-sea fish visual photopigment was then used to carry out molecular dynamics simulations and analysis, as previously described [33, 34, 69]. Briefly, the software package GROMACS 2022.5 [71] was used for all 100 ns molecular dynamics simulations with the Charmm36m forcefield [72] in presence of an explicit bilayer consisting of 1-stearoyl-2-docosahexaenoylphosphatidylcholine (SDPC) lipids. Median values for Angle 3, Torsion 15, AUC of RMSF, and presence/absence of the DisulfideBridge were used as inputs for calculating λ_max_ using the statistical models described above.

### Retinal histology

Retinal morphology was assessed in a total of 20 individuals. For each, one eye was processed for histological analyses: *V. mabahiss* [pre-flexion larvae (*n*=2), flexion larvae (*n*=1), early-mid post-flexion larvae (*n*=6), late post-flexion larvae (*n*=2), adults (*n=*2)], *B. pterotum* [pre-flexion larvae (*n*=1), post-flexion larvae (*n*=2), adult (*n=*1)] and *M. mucronatus* [flexion larvae (*n*=1), post-flexion larvae (*n*=1), adult (*n*=1)]. Whole, enucleated eyes were post-fixed in 2.5% glutaraldehyde and 2% osmium tetroxide, progressively dehydrated in increasing concentrations of ethanol, infiltrated with EPON resin and polymerized at 60°C for 48 h. For light micrographs, 1 μm-thick radial sections were cut on a Leica ultramicrotome (Ultracut UC6) and stained with 0.5% toluidine blue. For transmission electron micrographs, radial 90 nm-thick sections were air-dried onto copper mesh grids, stained with lead citrate and uranyl acetate and imaged using a Hitachi HT 7700 transmission electron microscope. Differentiation between rod- and cone-like morphology was based on ultrastructural features (Table 1). Specifically, rod-like cells were characterised by long, cylindrical OS, closed OS disc membranes, and the absence of a paraboloid or oil droplet.

Conversely, photoreceptors were classified as cone-like if they had short, tapered OS and open, continuous OS disc membranes. The quality of the samples did not permit clear morphological observations of the synaptic terminals, so these were not considered.

