## Supplemental Information for "Deep-sea fish reveal alternative pathway for vertebrate visual development"

### Extended Data

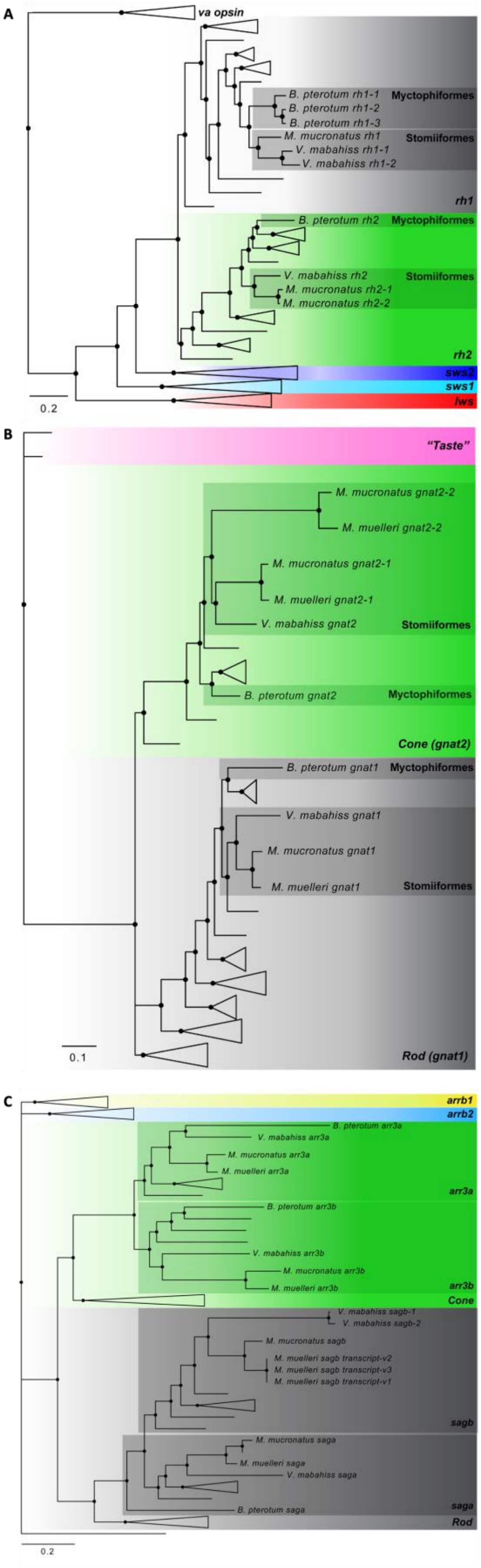

*Fig. S1. Gene phylogenies.* Vertebrate opsin (**A**), transducin (**B**) and arrestin (**C**) gene trees. The position of genes expressed in *V. mabahiss, M. mucronatus* and *B. pterotum* are highlighted. Overlay colours delineate gene families or subfamilies. In all panels, green-cone-specific genes are coloured in green while rod-specific genes are coloured in grey. Black circles denote Bayesian posterior probabilities > 0.8. The scale bars denote substitutions per site. Detailed phylogenies are given in Fig. S2 – S4.

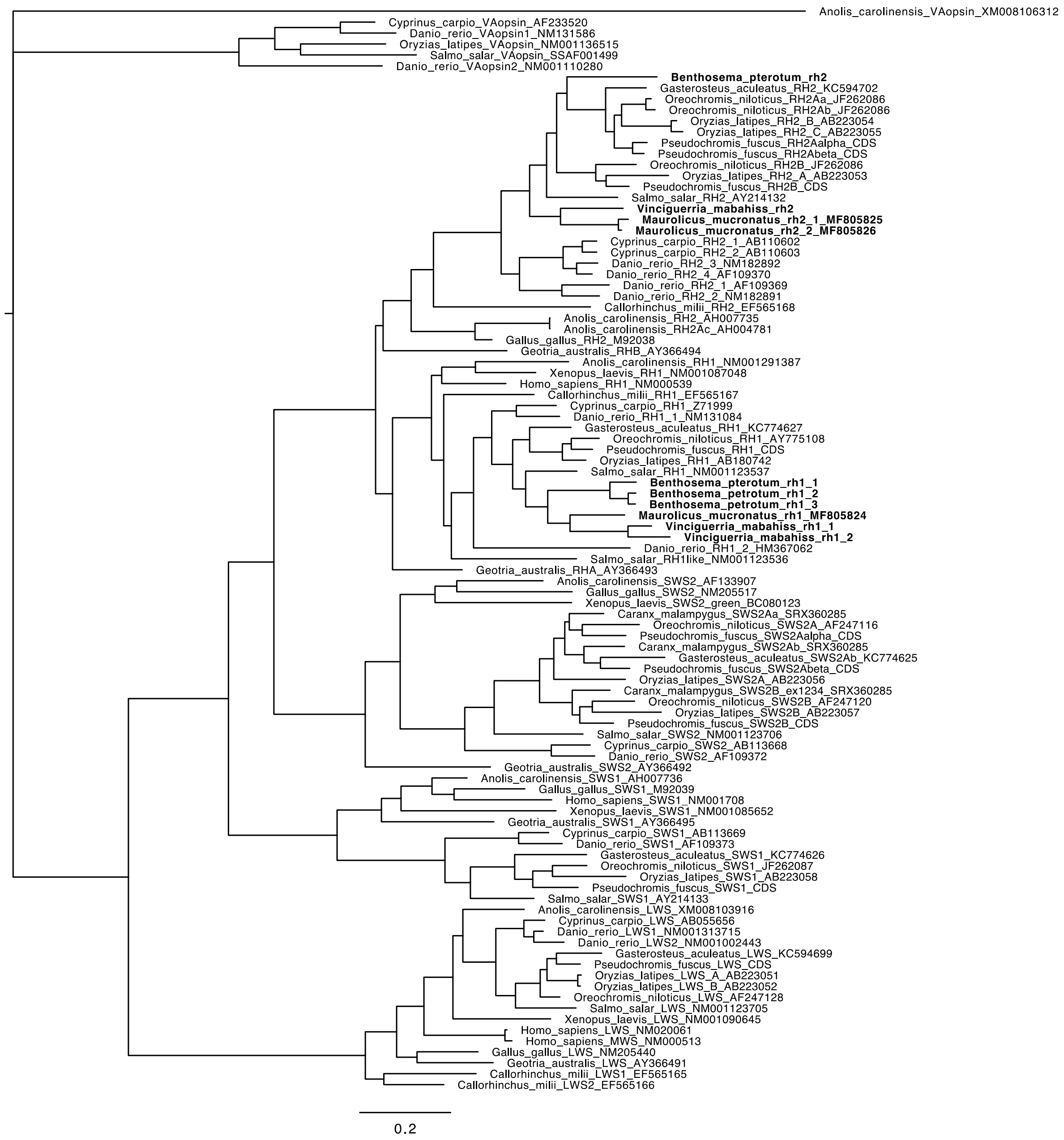


*Fig. S2. Expanded vertebrate opsin gene phylogeny.* Fully expanded opsin gene phylogeny. Placement of opsin genes extracted from *V. mabahiss, B. pterotum* and *M. mucronatus* are highlighted in bold.

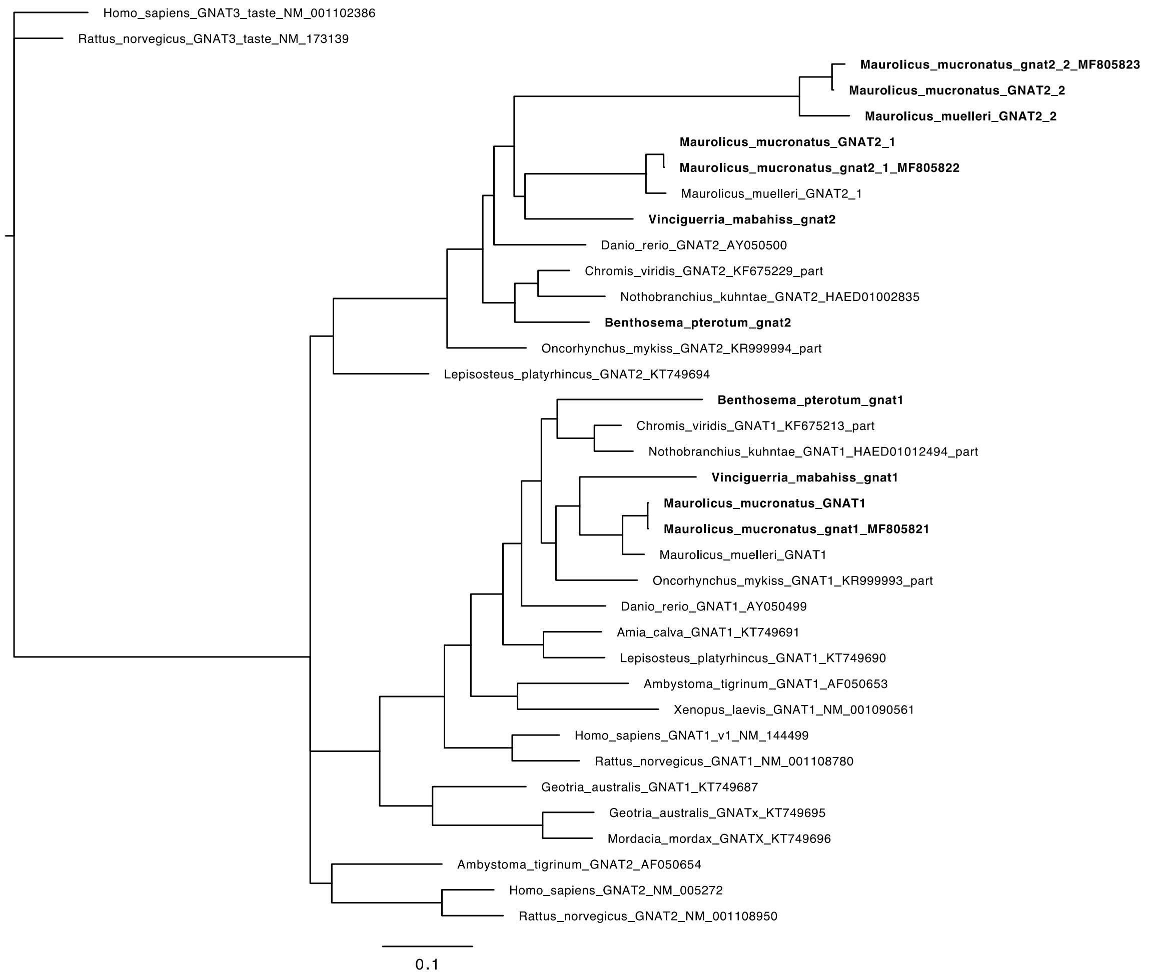


*Fig. S3. Expanded vertebrate transducin gene phylogeny.* Fully expanded transducin gene phylogeny. Placement of transducin genes extracted from *V. mabahiss*, *B. pterotum* and *M. mucronatus* are highlighted in bold.

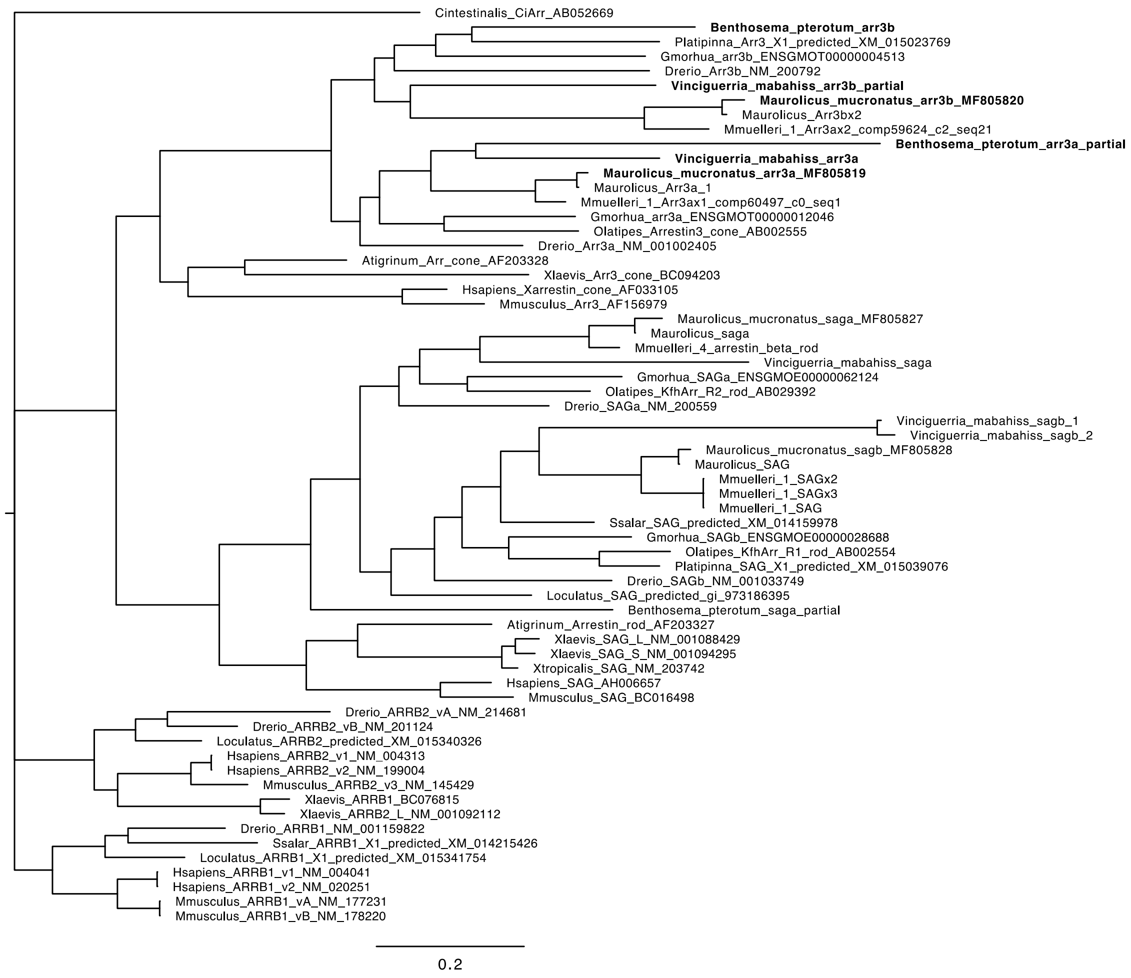


*Fig. S4. Expanded vertebrate arrestin gene phylogeny.* Fully expanded arrestin gene phylogeny. Placement of arrestin genes extracted from *V. mabahiss, B. pterotum* and *M. mucronatus* are highlighted in bold.

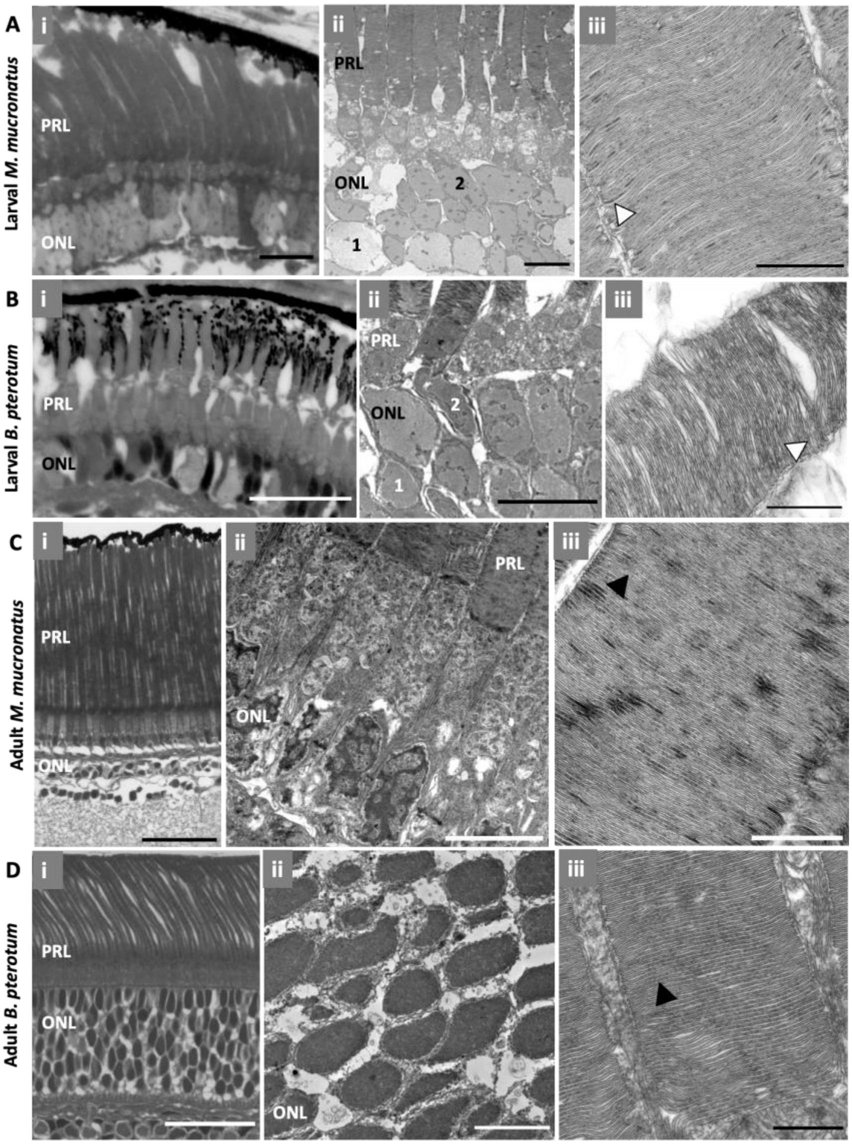


*Figure S5. Photoreceptor morphology in M. mucronatus and B. pterotum.* Representative light (**i**) and transmission electron (**ii, iii**) micrographs of the larval and adult photoreceptor layers of *M. mucronatus* (**A, C**), and *B. pterotum* (**B, D**), respectively. **(A – B)** In larvae, the retinae were dominated by rod-like photoreceptors, with long, cylindrical outer segments (**i**) and closed outer segments discs (**iii**; white arrow). Notably, all larvae had two morphological types of nuclei in the outer nuclear layer (ONL) characterised by lighter (type 1) or darker (type 2) chromatin staining (**ii**). **(C – D)** Like in larvae, adult retinae were also dominated by rod-like photoreceptors, with long, cylindrical outer segments (**i**), and closed outer segments discs (**iii**; black arrow). However, adults had only one type of nuclei in the ONL, which showed darker chromatin staining (**ii**). PRL, photoreceptor layer; ONL, outer nuclear layer. Scale bars: Ai, 10 µm; Bi, Di, 25 µm; Ci, 40 µm; A-Dii, 5 µm; Biii, Diii, 500 nm; Aiii, Ciii, 1 µm.

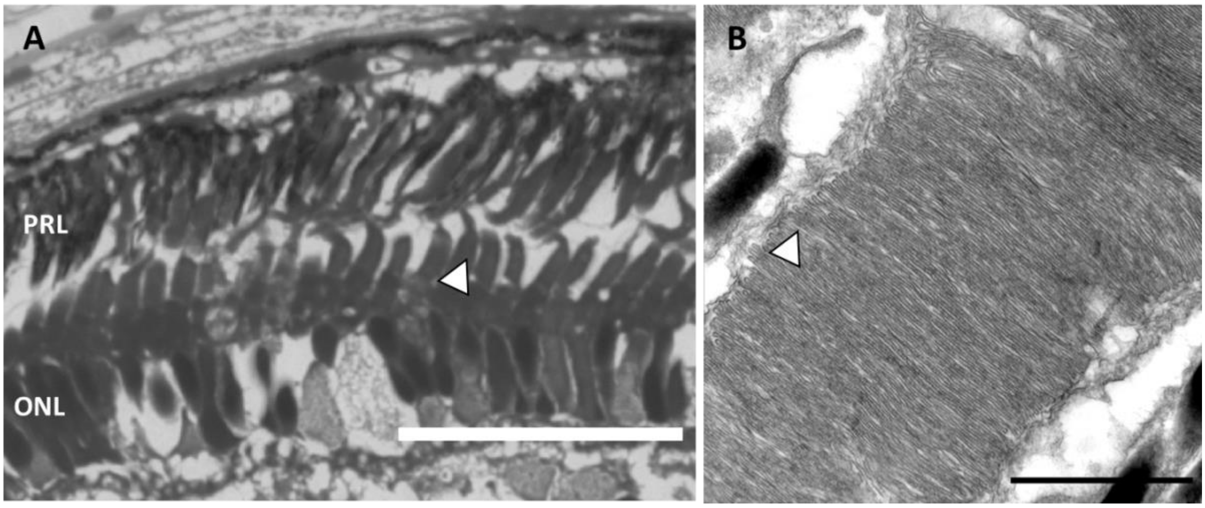


*Fig. S6. Morphology of cone-like photoreceptors in B. pterotum.* **A-B**. Micrographs of the retina in larval *B. pterotum*, showing overall morphology of a cone-like photoreceptor (arrow) within the photoreceptor layer (PRL) under light microscopy (**A**) and open (cone-like) outer segment discs (arrow) under TEM (**B**). ONL, outer nuclear layer. Scale bars: A, 25 µm; B, 1 µm.

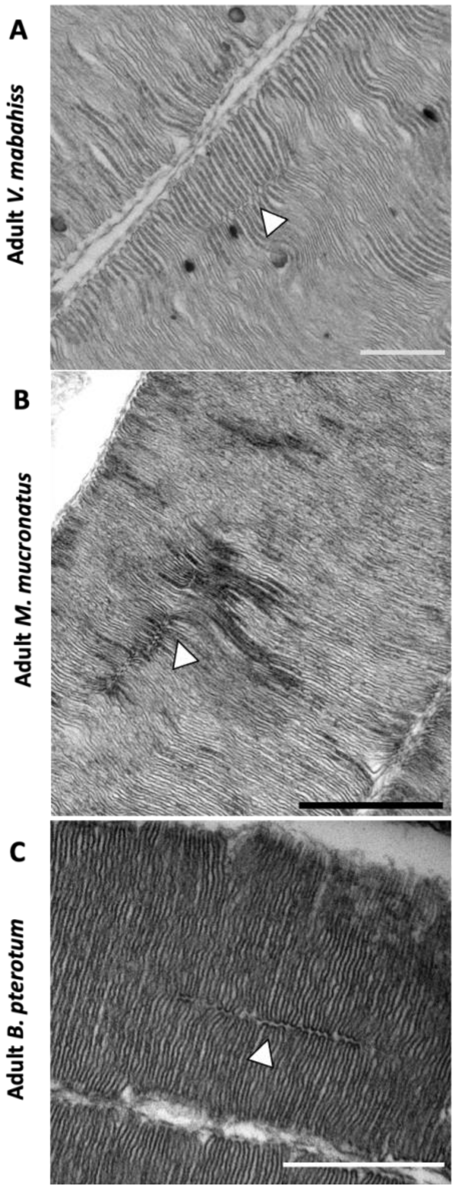


*Fig. S7. Incisures in rod-like photoreceptors of adult fishes.* Representative transmission electron micrographs of the photoreceptors of adult *V. mabahiss* (**A**), *M. mucronatus* (**B**), and *B. pterotum* (**C**). In adults of all species, the retina was dominated by rod-like photoreceptors, all of which had incisures (white arrow) in their outer segments. Scale bars: Div, Fiv, 500 nm; Eiv, 1 µm.

*Table S1. Details of animals used in study.* All fish were collected from the Saudi Arabian Red Sea. *Individual was in a mixed batch and exact specimen is unknown so a range is given for the standard length of all individuals in the batch.

| **Species** | **Life stage** | **SL (mm)** | **Analyses** |
|---|---|---|---|
| *V. mabahiss* | Preflexion | 4.8 | TEM, Light microscopy |
| | | 5.3 | TEM, Light microscopy |
| | | 3.5 | RNAseq |
| | | 3.7 | RNAseq |
| | | 4.2 | RNAseq |
| | Flexion | 5.5 | TEM, Light microscopy |
| | Early postflexion | 8.0 | Light microscopy |
| | | 5.6 | TEM, Light microscopy |
| | | 6.4 | TEM, Light microscopy |
| | | 7.2 | TEM, Light microscopy |
| | | 7.0 | RNAseq |
| | | 5.9 | RNAseq |
| | | 7.6 | RNAseq |
| | Mid postflexion | 11.0 | Light microscopy |
| | | 11.2 | TEM, Light microscopy |
| | | 11.0 | RNAseq |
| | Late postflexion | 12.2 | TEM, Light microscopy |
| | | 11.7 | Light microscopy |
| | | 12.6 | RNAseq |
| | Adult | 18.0 | TEM, Light microscopy |
| | | 14.6 | TEM, Light microscopy |
| | | 19.3 | RNAseq |
| | | 17.2 | RNAseq |
| | | 16.0 | RNAseq |
| | | 15.5 | RNAseq |
| | | 15.1 | RNAseq |
| *B. pterotum* | Preflexion | 3.7 | TEM, Light microscopy |
| | | 2.5 | RNAseq |
| | Postflexion | 4.8 | TEM, Light microscopy |
| | | 5.0 | TEM, Light microscopy |
| | | 5.8 | RNAseq |
| | Adult | 31.7 | TEM, Light microscopy |
| | | 25.3 | RNAseq |
| | | 26.1 | RNAseq |
| | | 28.1 | RNAseq |
| | | 23.0 | RNAseq |
| | | 22.0 | RNAseq |
| *M. mucronatus* | Flexion | 5.4 | TEM, Light microscopy |
| | | 4.0 | RNAseq |
| | Postflexion | 7.8 | TEM, Light microscopy |
| | | 7.3 – 9.4* | RNAseq |

| | | | |
|---|---|---|---|
| | Adult | 17.1 | TEM, Light microscopy |
| | | 23.4 | RNAseq |
| | | 22.3 | RNAseq |
| | | 22.4 | RNAseq |
| | | 21.5 | RNAseq |
| | | 21.2 | RNAseq |

*Table S2. Opsin gene expression.* Per-specimen mapped reads, gene lengths, gene length-adjusted opsin gene expression and proportional opsin gene expression values (as % of total opsin gene expression).

| Species | Stage | ID | Gene | Reads mapped | Gene length (bp) | Gene length-adjusted expression | Expression (%) |
|---|---|---|---|---|---|---|---|
| ***V. mabahiss*** | PreF | 1 | *rh2* | 7454 | 1044 | 7.14 | 100.00 |
| | | 2 | *rh2* | 15766 | 1044 | 15.10 | 100.00 |
| | | 3 | *rh2* | 31272 | 1044 | 29.95 | 100.00 |
| | Early PF | 1 | *rh2* | 13136 | 1044 | 12.58 | 100.00 |
| | | 2 | *rh2* | 18584 | 1044 | 17.80 | 100.00 |
| | | 3 | *rh2* | 29616 | 1044 | 28.37 | 100.00 |
| | Mid PF | 1 | *rh2* | 42074 | 1044 | 40.30 | 100.00 |
| | Late PF | 1 | *rh1-1* | 89486 | 915 | 97.80 | 100.00 |
| | | | *rh1-2* | 185202 | 915 | 202.41 | 100.00 |
| | Adult | 1 | *rh1-1* | 500449 | 915 | 546.94 | 60.26 |
| | | | *rh1-2* | 330013 | 915 | 360.67 | 39.74 |
| | | 2 | *rh1-1* | 338477 | 915 | 369.92 | 58.67 |
| | | | *rh1-2* | 238435 | 915 | 260.58 | 41.33 |
| | | 3 | *rh1-1* | 341833 | 915 | 373.59 | 46.47 |
| | | | *rh1-2* | 393799 | 915 | 430.38 | 53.53 |
| | | 4 | *rh1-1* | 338477 | 915 | 369.92 | 58.67 |
| | | | *rh1-2* | 238435 | 915 | 260.58 | 41.33 |
| | | 5 | *rh1-1* | 500498 | 915 | 546.99 | 60.26 |
| | | | *rh1-2* | 330062 | 915 | 360.72 | 39.74 |
| ***B. pterotum*** | PreF | 1 | *rh1* | 35 | 1047 | 0.03 | 0.21 |
| | | | *rh2* | 17005 | 1050 | 16.20 | 99.79 |
| | PF | 1 | *rh1* | 283 | 1047 | 0.27 | 3.00 |
| | | | *rh2* | 9187 | 1050 | 8.75 | 97.00 |
| | Adult | 1 | *rh1-1* | 170413 | 1050 | 162.30 | 23.99 |
| | | | *rh1-2* | 540017 | 1050 | 514.30 | 76.01 |
| | | 2 | *rh1-1* | 121126 | 1050 | 115.36 | 20.56 |
| | | | *rh1-2* | 468094 | 1050 | 445.80 | 79.44 |
| | | 3 | *rh1-1* | 268320 | 1050 | 255.54 | 28.09 |
| | | | *rh1-2* | 687028 | 1050 | 654.31 | 71.91 |
| | | 4 | *rh1-1* | 155649 | 1050 | 148.24 | 21.87 |
| | | | *rh1-2* | 556111 | 1050 | 529.63 | 78.13 |
| | | 5 | *rh1-1* | 72873 | 1050 | 69.40 | 15.47 |
| | | | *rh1-2* | 398085 | 1050 | 379.13 | 84.53 |
| ***M. mucronatus*** | F | 1 | *rh2-1* | 90524 | 1047 | 86.46 | 53.71 |
| | | | *rh2-2* | 78030 | 1047 | 74.53 | 46.29 |

| | | | | | | |
|---|---|---|---|---|---|---|
| | | *rh1* | 0 | 1074 | 0.00 | 0.00 |
| PF | 1 | *rh2-1* | 390445 | 1047 | 372.92 | 47.60 |
| | | *rh2-2* | 428539 | 1047 | 409.30 | 52.25 |
| | | *rh1* | 1263 | 1074 | 1.18 | 0.15 |
| Adult | 1 | *rh1* | 2544 | 1074 | 2.37 | 0.38 |
| | | *rh2-1* | 97713 | 282 | 346.50 | 55.69 |
| | | *rh2-2* | 79531 | 291 | 273.30 | 43.93 |
| | 2 | *rh1* | 1109 | 1074 | 1.03 | 0.46 |
| | | *rh2-1* | 34687 | 282 | 123.00 | 54.50 |
| | | *rh2-2* | 29581 | 291 | 101.65 | 45.04 |
| | 3 | *rh1* | 4197 | 1074 | 3.91 | 0.41 |
| | | *rh2-1* | 158531 | 282 | 562.17 | 59.47 |
| | | *rh2-2* | 110361 | 291 | 379.25 | 40.12 |
| | 4 | *rh1* | 1203 | 1074 | 1.12 | 0.20 |
| | | *rh2-1* | 91573 | 282 | 324.73 | 58.35 |
| | | *rh2-2* | 67131 | 291 | 230.69 | 41.45 |
| | 5 | *rh1* | 4023 | 1074 | 3.75 | 0.43 |
| | | *rh2-1* | 140129 | 282 | 496.91 | 56.39 |
| | | *rh2-2* | 110723 | 291 | 380.49 | 43.18 |

*Table S3. Developmental transcription factor gene expression.* Per-specimen transcription factor gene expression (in $log_{10}$TPM) in the retinas of deep-sea fishes at different life stages.

| | otx5 | rorb | nr2e3 | nrl | thrb |
|---|---|---|---|---|---|
| **Preflexion *V. mabahiss*_1** | 5.46 | 5.61 | -1.00 | -1.00 | -1.00 |
| **Preflexion *V. mabahiss*_2** | 5.68 | 5.50 | 4.56 | -1.00 | -1.00 |
| **Preflexion *V. mabahiss*_3** | 5.80 | 5.27 | 4.58 | -1.00 | -1.00 |
| **Early-mid postflexion *V. mabahiss*_1** | 5.72 | 5.35 | 4.51 | -1.00 | -1.00 |
| **Early-mid postflexion *V. mabahiss*_2** | 5.85 | 4.98 | -1.00 | -1.00 | -1.00 |
| **Early-mid postflexion *V. mabahiss*_3** | 5.59 | 5.58 | 4.60 | -1.00 | -1.00 |
| **Early-mid postflexion *V. mabahiss*_4** | 5.95 | 4.09 | 5.02 | -1.00 | -1.00 |
| **Late postflexion *V. mabahiss*** | 5.89 | 5.05 | 3.87 | 4.74 | 4.26 |
| **Adult *V. mabahiss*_1** | 5.62 | 5.33 | 4.07 | 4.83 | 5.05 |
| **Adult *V. mabahiss*_2** | 5.67 | 5.25 | 3.94 | 5.02 | 4.99 |
| **Adult *V. mabahiss*_3** | 5.68 | 5.25 | 3.86 | 4.96 | 4.98 |
| **Adult *V. mabahiss*_4** | 5.67 | 5.25 | 3.96 | 5.02 | 4.99 |
| **Adult *V. mabahiss*_5** | 5.62 | 5.32 | 4.00 | 4.83 | 5.05 |
| **Preflexion *B. pterotum*** | 5.84 | 4.89 | 5.38 | -1.00 | -1.00 |
| **Postflexion *B. pterotum*** | 5.54 | 5.81 | -1.00 | -1.00 | -1.00 |
| **Adult *B. pterotum*_1** | 5.53 | 4.30 | 3.90 | 5.59 | 5.02 |
| **Adult *B. pterotum*_2** | 5.78 | 4.37 | 3.97 | 4.92 | 4.96 |
| **Adult *B. pterotum*_3** | 5.77 | 4.33 | 4.08 | 4.91 | 5.01 |
| **Adult *B. pterotum*_4** | 5.64 | 4.40 | 3.70 | 5.40 | 5.00 |
| **Adult *B. pterotum*_5** | 5.66 | 4.42 | 3.62 | 5.33 | 4.99 |
| **Flexion *M. mucronatus*** | 5.60 | 5.42 | -1.00 | 5.11 | -1.00 |
| **Postflexion *M. mucronatus*** | 5.78 | 4.76 | -1.00 | 4.55 | -1.00 |
| **Adult *M. mucronatus*_1** | 5.92 | 4.90 | -1.00 | 4.48 | -1.00 |
| **Adult *M. mucronatus*_2** | 5.81 | 5.33 | -1.00 | 4.60 | -1.00 |
| **Adult *M. mucronatus*_3** | 5.91 | 5.04 | -1.00 | 4.38 | -1.00 |
| **Adult *M. mucronatus*_4** | 5.81 | 5.32 | -1.00 | 4.59 | -1.00 |
| **Adult *M. mucronatus*_5** | 5.93 | 4.93 | -1.00 | 4.45 | -1.00 |
